# Genomic Analysis of Emerging Florfenicol-Resistant *Campylobacter coli* Isolated from the Intestinal Cecal Contents of Cattle in the United States

**DOI:** 10.1101/604645

**Authors:** Shaohua Zhao, Sampa Mukherjee, Chih-Hao Hsu, Shenia Young, Cong Li, Heather Tate, Cesar A. Morales, Jovita Haro, Sutawee Thitaram, Glenn E. Tillman, Uday Dessai, Patrick McDermott

## Abstract

**Objective:** Genomic analyses were performed on florfenicol resistant (FFN^R^) *Campylobacter coli* (*C. coli*) isolated from cattle and the *cfr*(C) gene-associated multi-drug resistance (MDR) plasmid was characterized.

**Methods:** Sixteen FFN^R^ *C. coli* isolates recovered between 2013-2018 from beef cattle were sequenced using MiSeq. Genomes and plasmids were closed for three of the isolates using the PacBio^®^ system. Single nucleotide polymorphisms (SNPs) across the genome and the structures of MDR plasmids were investigated. Conjugation experiments were performed to determine the transferability of *cfr*(C) associated MDR plasmids. The spectrum of resistance encoded by the *cfr*(C) gene was further investigated by agar dilution antimicrobial susceptibility testing.

**Results:** All 16 FFN^R^ isolates were MDR and exhibited co-resistance to ciprofloxacin, nalidixic acid, clindamycin and tetracycline. All isolates shared the same resistance genotype, carrying *aph(3’)-III, hph, ∆aadE* (truncated), *bla*_OXA-61_*, cfr*(C), and *tet*(O) genes plus a mutation of GyrA T86I. The *cfr*(C), *aph(3’)-III, hph ∆aadE*, and *tet*(O) genes were co-located on transferable MDR plasmids with size 48-50 kb. These plasmids showed high sequence homology with the pTet plasmid, and carried several *Campylobacter* virulence genes, including *virB2, virB4, virB5, VirB6, virB7, virB8, virb9, virB10, virB11 and virD4*. The *cfr*(C) gene conferred resistance to florfenicol (8-32 µg/ml), clindamycin (512-1,024 µg/ml), linezolid (128-512 µg/ml), and tiamulin (1,024 µg/ml). Phylogenetic analysis showed SNP differences ranging from 11-2,248 among the 16 isolates.

**Conclusions:** The results showed that the *cfr*(C) gene located in the conjugative pTet MDR/virulence plasmid is present in diverse strains, where it confers high levels of resistance to several antimicrobials, including linezolid, a critical drug for treating Gram positive bacterial infections in humans. This study highlights the power of genomic antimicrobial resistance surveillance to uncover the intricacies of transmissible co-resistance and provides information that is needed for accurate risk assessment and mitigation strategies.

## Introduction

*Campylobacter* is one of the leading bacterial causes of foodborne illness in the United States. Human infections are associated mainly with raw or undercooked chicken meat, but other sources such as beef, pork, lamb, water and seafood also have been associated with *Campylobacter* infections [1]. Antimicrobial resistance in *Campylobacter* is a public health concern [2–4]. In 2013, the Center for Disease Control and Prevention (CDC) classified drug-resistant *Campylobacter* as a serious threat in the United States (http://www.cdc.gov/drugresistance/threat-report-2013). The use of antimicrobials in animals, and the potential contribution to generating resistance in foodborne bacteria, has been an important public health issue for many years. The US National Antimicrobial Resistance Monitoring System (NARMS) was launched in 1996 to track changes in antimicrobial resistance in foodborne pathogens, including *Campylobacter*, isolated from food animals, retail meats, and humans. Currently, nine antimicrobials belonging to seven classes are included in the NARMS *Campylobacter* testing panel.

Florfenicol belongs to a class of phenicol antimicrobials that is approved in the US for treatment of bovine and swine respiratory infections [5, 6]. Since 2004, NARMS has monitored resistance to florfenicol in *Campylobacter* and resistance has been very rare in human and food isolates, although resistance has only been monitored in cecal samples since 2013. The first florfenicol resistant (FFN^R^) *Campylobacter coli* strains were detected in 2013 in cecal samples from beef cattle accounting for 1.6% of beef isolates tested (n=128) and increased to 4.4% resistant in 2014 (n=180). No FFN^R^ *C. coli* were detected in 2015 (n=181), but it reappeared in 2016 (n=200) and 2017 (n=239) accounting for 2% and 0.8% of resistance in beef *C. coli* isolates tested, respectively. All FFN^R^ *Campylobacter* isolates were *C. coli* recovered from cecal contents collected from beef cattle post-slaughter prior to any interventions steps. Antimicrobial susceptibility testing (AST) showed that all FFN^R^ *C. coli* were multidrug resistant (MDR) and showed resistance to five of nine antimicrobials tested, including resistance to ciprofloxacin, clindamycin, florfenicol, nalidixic acid, and tetracycline.

The *cfr*(C) gene was first reported in 2017 by Tang et al. and was shown to be responsible for florfenicol resistance in *C. coli* [7]. The *cfr*(C) gene encodes a protein that shares 55.1% and 54.9% amino acid identity with Cfr and Cfr(B), respectively [7]. The *cfr* gene was first detected in *Staphylococcus sciuri* isolated from a bovine origin in 2000 [8], and was later detected in *Enterococcus faecium* (*E. faecium*) isolated from a human bloodstream infection in 2015 [9]. The *cfr*(B) gene also has been detected in *Clostridium difficile* and *E. faecium* from humans [10, 11]. Although the three *cfr* alleles show high sequence diversity, all of them confer resistance to five chemically unrelated antimicrobial classes, including phenicols, lincosamides, oxazolidinones, pleuromutilins and streptogramins (PhLOPS_A_ phenotype) [7, 12]. Previous studies showed that *cfr, cfr*(B) and *cfr*(C) genes are located on various plasmids [7, 10, 11].

In the original study by Tang et al. [7], all FFN^R^ *C. coli* were isolated from cattle, with a 10% prevalence rate, but no FFN^R^ *Campylobacter jejuni* were detected. Their study showed that the *cfr*(C) gene located in the conjugative MDR plasmid also carried several other resistance genes including *tet*(O), *hph*, and *aphA-3*, which conferred resistance to tetracycline, hygromycin, and kanamycin, respectively. The plasmid also carried a truncated *aadE* gene. Pulsed-field gel electrophoresis (PFGE) and multi-locus sequence typing (MLST) analysis showed that the increasing prevalence of *cfr*(C) in *C. coli* is due to clonal expansion [7]. To further understand the mechanism of FFN^R^, and the genetic context of its spread over time, we performed whole genome sequencing (WGS) analyses on 16 FFN^R^ *C. coli* isolates recovered between 2013-2018 to identify the resistance genotype and characterize FFN^R^ MDR plasmids.

## Materials and Methods

### Bacterial strains

All 16 FFN^R^ *C. coli* isolates were recovered from beef cattle cecal contents between 2013-2018 as part of the US NARMS program (Table 1). The isolates were obtained by the U. S. Department of Agriculture Food Safety and Inspection Service (USDA FSIS). The isolates were grown on sheep blood agar plates (Thermo Fisher Scientific, Remel, Lenexa, KS) at 42°C under microaerobic conditions (85% N2, 10% CO2, and 5% O2). WGS data of the FFN^R^ *C. coli* isolates recovered from cattle cecal contents at slaughter were generated by FSIS or FDA. One florfenicol susceptible (FFN^S^), erythromycin resistant (ERY^R^) *C. jejuni* isolate (N18880) recovered from retail chicken was used as the recipient strain for conjugation experiments. The resistance phenotypes of all 16 isolates were previously determined by the broth-micro dilution method using the NARMS *Campylobacter* panel [13].

**Table 1:**
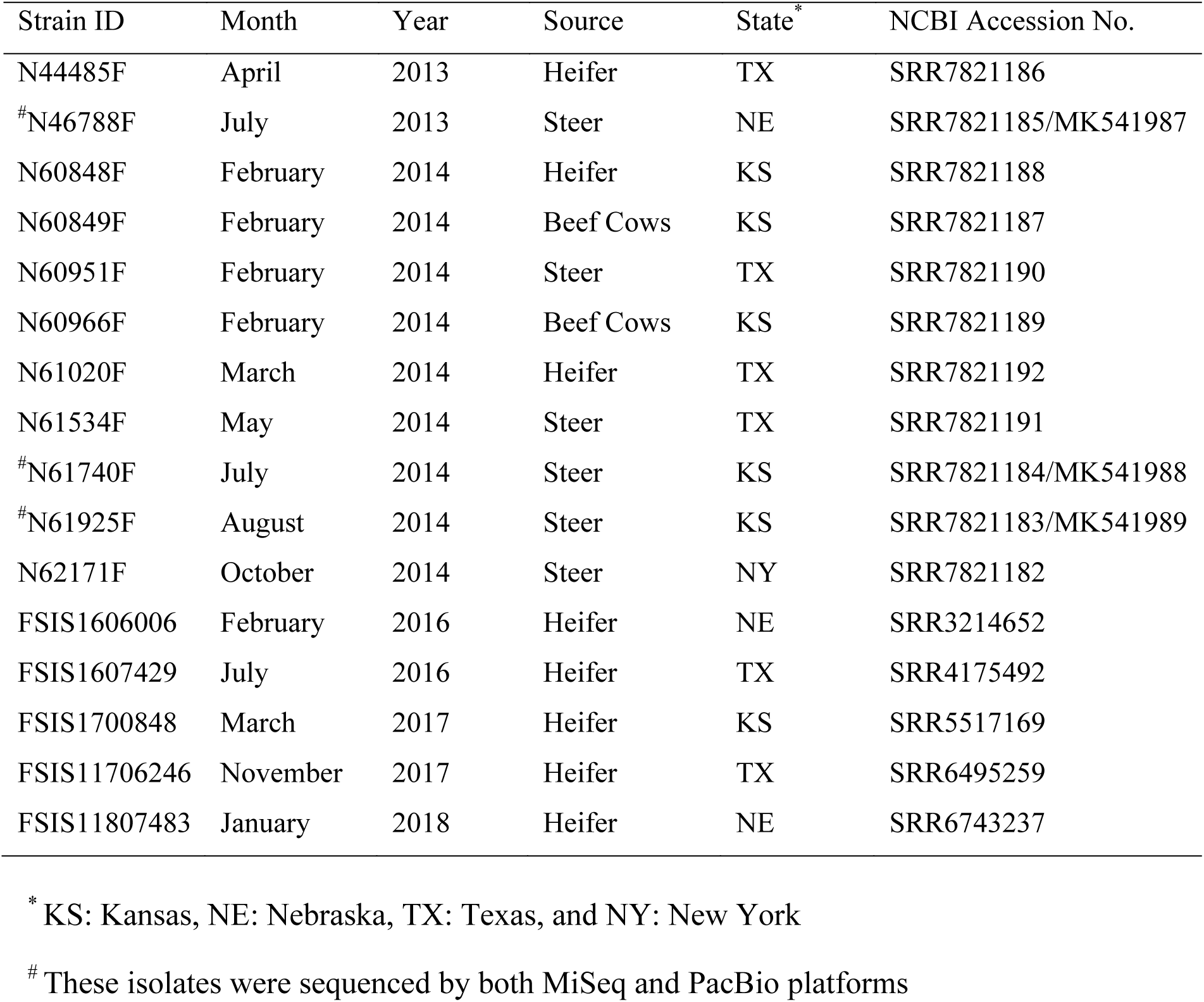
Florfenicol Resistant *Campylobacter coli* Strains in this Study

### Genome sequencing, assembly and annotation

Genomic DNA was extracted with QIAamp 96 DNA QIA cube HT kit (Qiagen, Gaithersburg, MD, USA) on an automated high-throughput DNA extraction machine (QIAcube HT, source) per the manufacturer’s instructions. WGS was performed on the Illumina MiSeq platform using v3 reagent kits (Illumina, San Diego, CA, USA) and the 2×300 paired end option. Assembly was performed de novo for each isolate using CLC Genomics Workbench version 8.0 (CLC bio, Aarhus, Denmark). Three isolates (N46788F, N61740F, and N61925F) were selected to close the genomes and plasmids using the Pacific Biosciences (PacBio) RS II Sequencer (PacBio^®^, Menlo Park, CA. USA). The continuous-long-reads were assembled by PacBio Hierarchical Genome Assembly Process (HGAP3.0) program. Genomes were annotated using the RAST annotation server (http://rast.nmpdr.org/). Among the 16 FFN^R^ *C. coli* isolates sequenced on the MiSeq, there was a median of 70 contigs (ranging from 23-573) and 103 fold coverage (ranging from 26-138) per genome.

### Identification of antimicrobial resistance genotypes

Antimicrobial resistance genes were identified using Perl scripts to perform local BLAST with ResFinder (https://cge.cbs.dtu.dk/services/ResFinder/) with at least 85% nucleotide identity and 50% sequence length to known resistance gene sequences. Sequences showing less than 100% identity and/or sequence length were examined by additional BLAST analysis to identify the appropriate resistance genes. Mutations in *gyrA* gene were identified using an in-house pipeline[14].

### Plasmid Analysis

The pTet (81-176) plasmid sequences were downloaded from NCBI GenBank under accession number AY394561. Our *cfr*(C) MDR plasmid sequences were blasted against the pTet plasmid sequences to determine sequence homology. The pTet plasmid and our *cfr*(C) MDR plasmids were also annotated using the RAST annotation server to compare the annotated genes among these plasmids.

### Whole genome phylogenetic analysis

The single nucleotide polymorphism (SNP) analysis of 16 FFN^R^ *C. coli* isolates was performed using the Food and Drug Administration Center for Food Safety and Applied Nutrition (CFSAN)-SNP-Pipeline (http://snp-pipeline.readthedocs.io/en/latest/). The complete genome of *C. coli* strain MG1116 (NCBI accession number CP017868) was used as a reference genome. VarScan [15] was used to detect SNPs. Plasmid sequences were excluded in the SNP analysis. SNP redundancy by linkage disequilibrium (LD) was reduced and the phylogenetic tree was constructed with the maximum likelihood algorithm using the SNPhylo package [16].

### Conjugation and antimicrobial susceptibility testing

Two FNN^R^ *C. coli* isolates (N61740F and N61925F) that carried *cfr*(C) gene were used as donor strains and one florfenicol susceptible (FNN^S^), ERY^R^ *C. jejuni* (N18880) was used as recipient strain (Table 2) in agar plate mating experiments as previously described by Chen [17]. Briefly, to prepare donor and recipient strains, one loopful of bacteria grown overnight on a sheep blood agar plate was re-suspended in 200 µl LB broth; 10µl of each donor strain was spotted separately on top of seven 10µl spots of recipient strain on a fresh sheep blood agar plate. Plates were incubated overnight at 42⁰C under microaerobic conditions. Each co-culture was scraped from the plate and re-suspended in 500 µl LB broth. 100µl each of 1:10 and 1:100 dilutions of the re-suspension were plated on agar plates supplemented with appropriate selective agents. Florfenicol (8µg/ml) and erythromycin (16 µg/ml) were used as selecting markers for the conjugation experiment. Successful transconjugants were confirmed by the comparing resistant phenotypes of donors, recipient and transconjugants using the Sensititre^TM^ automated antimicrobial susceptibility system in accordance with the manufacturer’s instructions (ThermoFisher Scientific, Trek Diagnostics, Cleveland, OH) using the NARMS *Campylobacter* panel (catalog#: CAMPY). Nine antimicrobial agents were tested, including azithromycin (AZI), ciprofloxacin (CIP), clindamycin (CLI), erythromycin (ERY), florfenicol (FFN), gentamicin (GEN), nalidixic acid (NAL), telithromycin (TEL), and tetracycline (TET). *C. jejuni* ATCC 33560 was used as the quality control organism according to guidelines of the Clinical and Laboratory Standards Institute (CLSI). Interpretation of susceptibility testing results was based on the European Committee on Antimicrobial Susceptibility Testing (EUCAST) epidemiological cutoff values (http://www.eucast.org/). PCR analysis was used to confirm that the transconjugants were *C. jejuni* [18].

**Table 2:** Antimicrobial Susceptibility of Donors, Recipients, and the Transconjugants.

| CVM# | MICs of Antimicrobials<br>Micro-broth Dilution (µg/ml) |  |  |  |  |  |  |  |  | MICs of Antimicrobials<br>Agar Dilution (µg/ml) |  |  |  |
| --- | --- | --- | --- | --- | --- | --- | --- | --- | --- | --- | --- | --- | --- |
|  | AZI | CIP | CLI | ERY | FFN | GEN | NAL | TEL | TET | LZD | TIA | CLI | Description |
| N61740F | 0.12 | 16 | >16 | 2 | 32 | 0.5 | >64 | 2 | >64 | 256 | 1024 | 1024 | Donor |
| N61925F | 0.12 | 16 | >16 | 2 | 16 | 0.5 | >64 | 2 | 64 | 256 | 1024 | 1024 | Donor |
| TCN61740F | >64 | 0.12 | >16 | 64 | 16 | 0.5 | ≤4 | 8 | 64 | 512 | 1024 | 1024 | Transconjugant |
| TCN61925F | >64 | 0.12 | 16 | 64 | 8 | 0.5 | ≤4 | 8 | >64 | 128 | 1024 | 512 | Transconjugant |
| N18880R | >64 | 0.12 | 4 | >64 | 1 | 0.5 | ≤4 | 8 | 0.5 | 16 | 2 | 16 | Recipient |

AZI: Azithromycin, CIP: Ciprofloxacin, CLI: Clindamycin, ERY: Erythromycin, FFN:
Florfenicol, GEN: Gentamycin, NAL: Nalidixic Acid, TEL: Telithromycin, TET: Tetracycline,
LZD: Linezolid, TIA: Tiamulin

To measure the contribution of *cfr*(C) to linezolid (LZD), tiamulin (TIA), and clindamycin (CLI) resistance, MICs were measured for the donors, recipient and transconjugants by agar dilution as described previously after transmission of the *cfr*(C) containing plasmid [19, 20]. Briefly, agar plates were prepared with four drugs with concentrations ranges from 0.125 µg/ml to 1024 µg/ml for each antimicrobial. Minimum inhibitory concentrations (MIC) were determined based on CLSI guidelines [21] and were recorded as the lowest concentration of antimicrobial agent that completely inhibited the visible growth of the organism on the agar surface after incubation at 42⁰C for 24 hours.

## Results and Discussion

### Resistance phenotypes and genotypes

All 16 FFN^R^ isolates were MDR, and showed co-resistance to CIP, NAL, CLI and TET tested using NARMS *Campylobacter* panel. Additionally, they shared the same resistance genotype, carrying *aph(3’)-III, hph, ∆aadE, bla*_OXA-61_, *cfr*(C), and *tet*(O) genes plus a same mutation of GyrA T86I (Figure 1). The FFN^R^ strain Tx40 reported by Tang was also MDR, and showed resistance to CIP, TET, CLI, FFN, LZD, TIA, chloramphenicol (CHL), and tedizolid (TED) [7].

**Figure 1.**
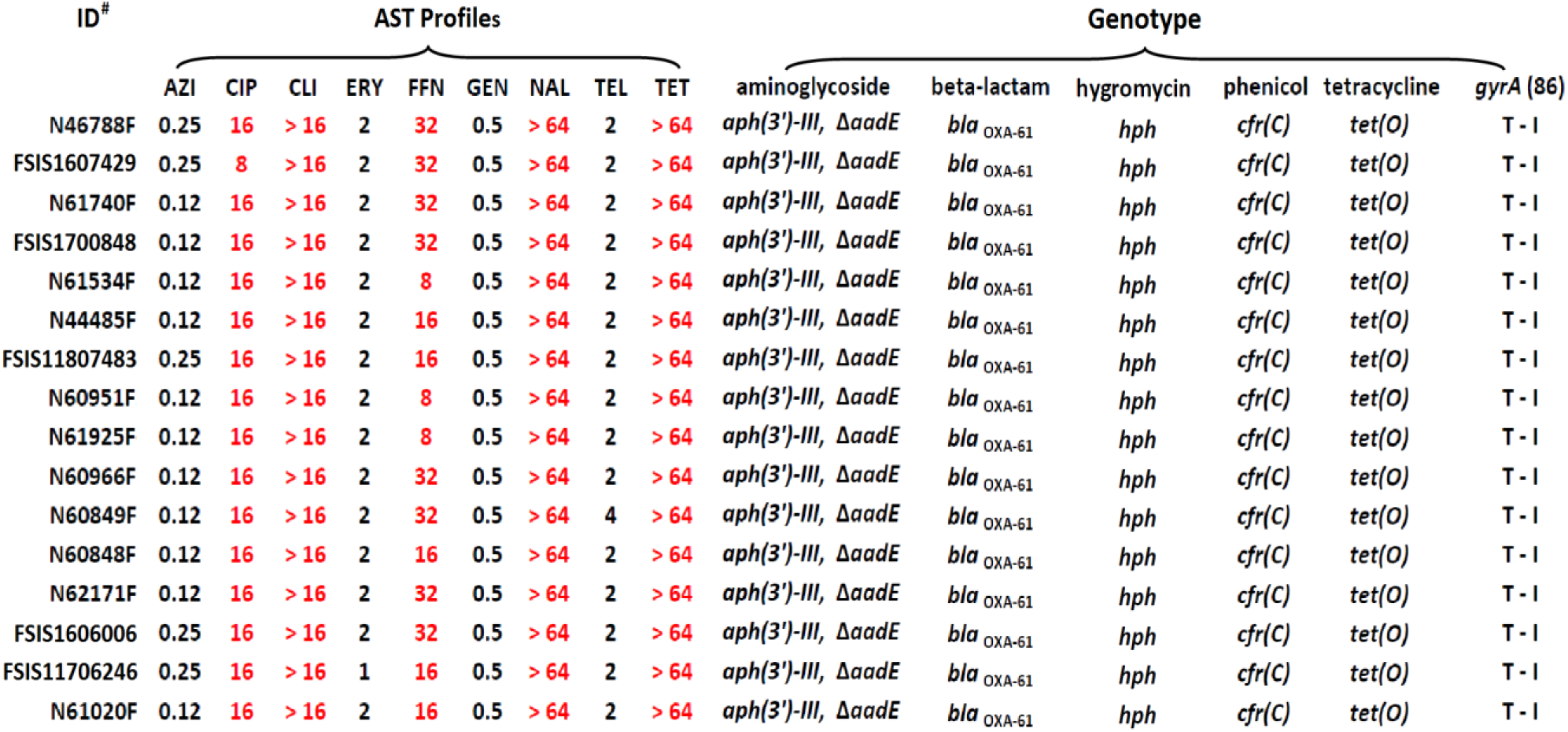
Antimicrobial Susceptibility Profiles and Resistance Genotypes of FFN^R^ *Campylobacter coli* Isolated Cattle

### Conjugation and *cfr*(C) co-resistance to other antimicrobials

Two FFN^R^ *C. coli* strains, N61740F and N61925F carrying *cfr*(C) gene were used as donors for the conjugation experiment. The results showed that the *cfr*(C) gene was successfully transferred to a FFN^S^ *C. jejuni* strain (N18880) based on species confirmation by PCR and the AST profiles of transconjugants. Two transconjugants (TCN61740F and TCN61925F) showed increasing MICs ≥4 fold and ≥8 fold for CLI and FFN, respectively, as compared to the N18880 parent recipient strain (Table 2). Agar dilution antimicrobial sensitivity testing determined that the MICs of CLI, LZD, and TIA increased 32-64, 8-32, and 512-fold, respectively, in two transconjugants as compared with the parent recipient strain (Table 2). Similar findings were reported from Tang’s study [7]. Their results showed that the MICs of transconjugant (JL272/pTx40) or cloning/transformant [*C.jejuni*11168/pRY108-*cf*r(C)] were 32, >16 and >128 fold increase for CLI, LZD, and TIA, respectively, as compared with their parent strains [7]. Both studies showed that the genetic background of recipient/parent cells, and different susceptibility testing methods could result in variations of MIC.

### MDR virulence plasmids

Three plasmids from strains N61925, N61740 and N46788F were closed using the PacBio long sequencing platform. Two plasmids, pN61925 and pN61740, were identical size, 48,049 bp with 99.9% sequence identity. The third plasmid, pN46788F, was 50,413 bp which has >91% sequence identity with pN61925 and pN61740. All three plasmids were annotated to encode similar 55 ORFs, including 22 encoding known function proteins and 23 hypothetical proteins (Figure 2). Among the genes with known functions, four resistance genes, *tet*(O), *hph, aph(3’)-III*, *cfr*(C), and one truncated resistance gene, ∆*aadE*, plus ten virulence genes, *virB2, virB4* (two copies), *virB5, virB6, virB7*, *virB8, VirB9, VirB10* and *VirB11* were identified (Figure 2). The pN61925, pN61740 and pN46788 plasmid structure and gene organization were the same as the pTx40 plasmid reported by Tang et.al [7]. We also compared DNA sequences and overall gene organization between the pN61925 plasmid and the pTet 81-176 plasmid, the first reported case of pTet plasmid from *Campylobacter* [22]. The two plasmids shared about 41 kb sequence, and the only sequence difference is a region that encodes antimicrobial resistance genes. The pTet 81-176 plasmid only carried the *tet*(O) gene, whereas, the pN61925 carried five resistance genes [∆*aadE, hph, aph(3’)-III, cfr*(C) and *tet*(O)] in addition to *pcp* and *tolA* (Figure 2). A plasmid with a similar structure as pTet 81-176 was also identified from *C. coli* isolated from NARMS retail chicken in early 2011 [17]. The plasmid carried several antibiotic resistance genes including a novel gentamicin resistance gene [*aph(2’’)-Ig*].

**Figure 2.**
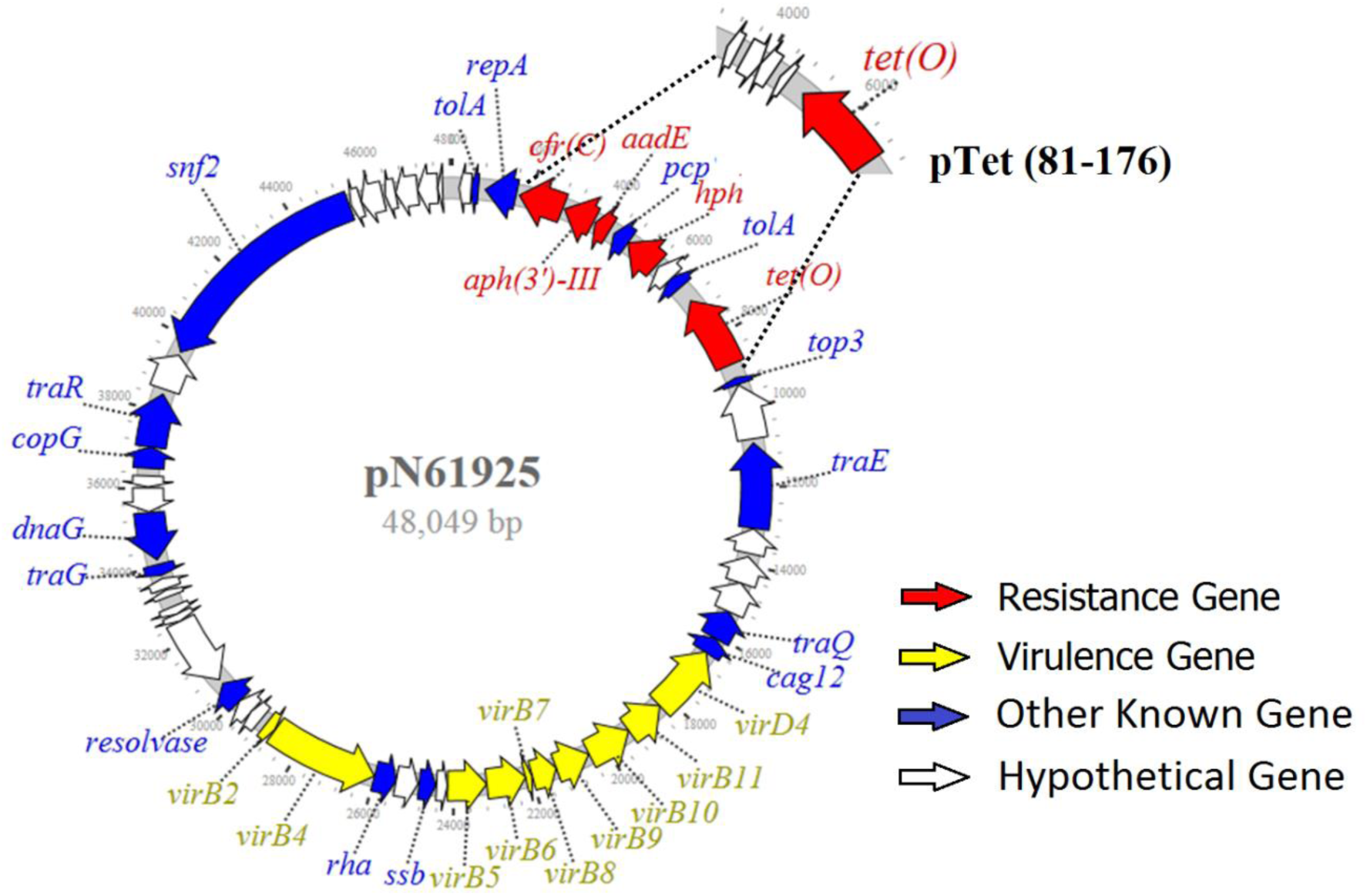
Structure of Multi-Drug Resistance/Virulence Plasmid from FFN^R^ *Campylobacter coli*

The virulence factors encoded by pTet and *cfr*(C) plasmids were previously reported [22–24]. These virulence factors are involved in bacterial pathogenesis, including adherence, invasion, motility, and immune evasion [23, 24]. Some of these virulence factors are involved with the structure of type IV secretion systems (T4SSs), which have been found in both Gram positive and Gram negative bacteria, including *Argobacterium tumefaciens, Bordetella pertussis*, several *Brucella species*, *C. jejuni*, *Helicobacter pylori*, and others [25, 26]. T4SSs comprise a class of diverse transporters and secrete a wide range of substrates, ranging from single proteins to protein-protein and protein-DNA complexes, which are required for virulence in many pathogens [25, 26]. More studies on *Campylobacter* pathogenesis associated with T4SSs and pTet MDR virulence plasmid are needed.

### Whole Genome SNP Analysis

The SNP tree of 16 FFN^R^ *C. coli* isolates showed that they were genetically diverse. The whole genome SNP (wgSNP) differences ranged from 11 to 2,248 despite some of isolates being collected in the same years from the same state (Figure 3). For example, all 9 isolates recovered in 2014 (five from KS, three from TX and one from NY) were scattered in all branches of the phylogenetic tree with a maximum 2,248 SNP differences even though some isolates are from the same states and they share the same resistance phenotype and genotypes (Figure 1). This is most likely due to all isolates containing the same MDR plasmid which carried the same resistance genes: *aph(3’)-III, ∆aadE, hph, cfr*(C) and *tet*(O). All isolates also carried a *bla*_OXA-61_ and had a mutation in GyrA T(86)I, which are responsible for beta-lactam and quinolone resistance phenotypes, respectively [14]. Previous studies showed that a *bla*_OXA-61_ is commonly present on the chromosome of *C. jejuni* and *C. coli* [27, 28].

**Figure 3.**
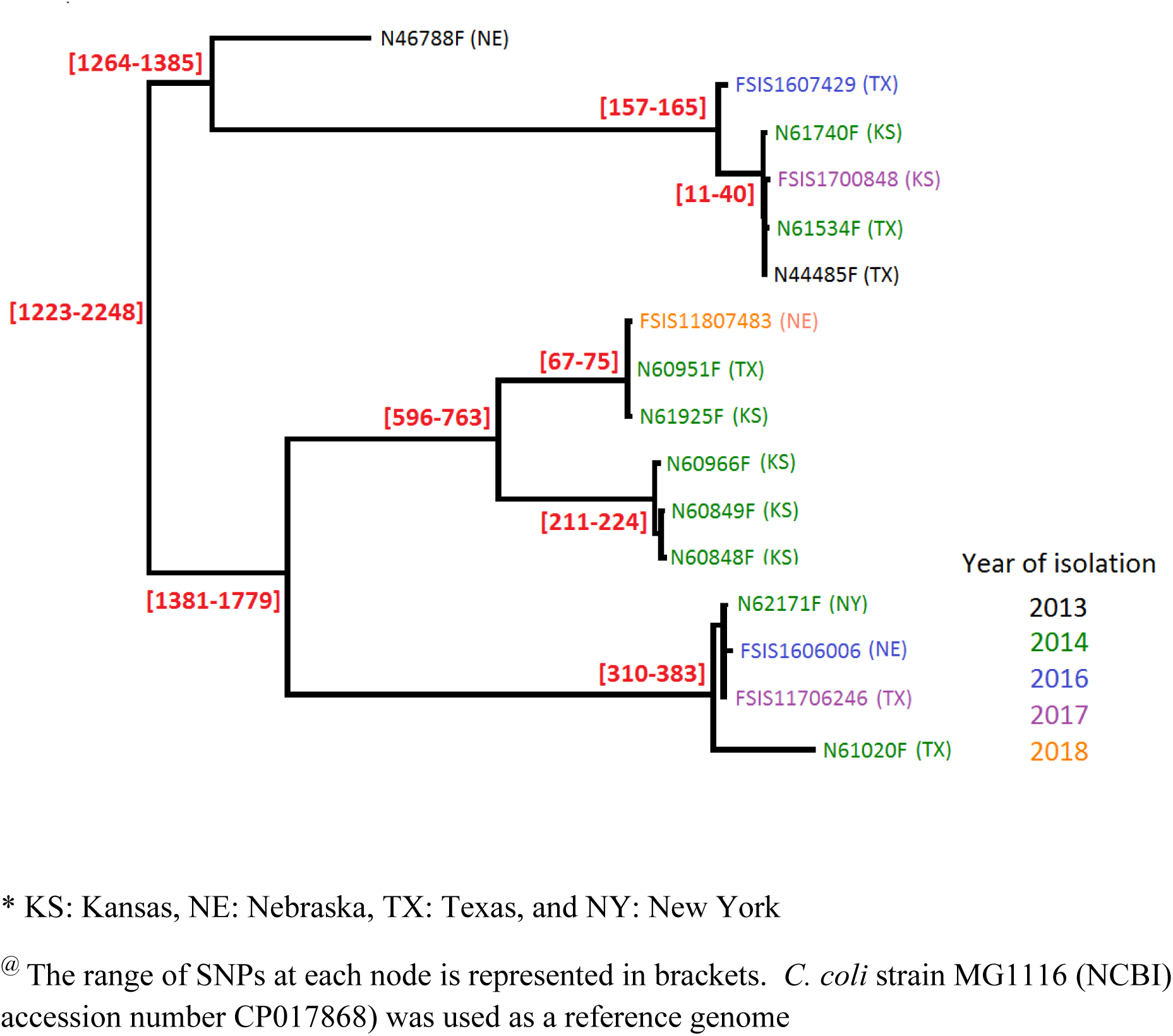
hqSNP Core-Genome Tree of FFN^R^ *Campylobacter coli* Strains Isolated from Cattle^@^.

Tang et.al characterized 34 *cfr*(C) positive *C. coli* isolates by PFGE and MLST and suggested that clonal expansion was involved in the spread of *cfr*(C) positive *C. coli* isolates in feedlot cattle in the US [7]. In contrast, our study suggests that spread of *cfr*(C) positive *C. coli* in cattle is most likely due to the spread of a MDR *cfr*(C) plasmid. The difference in these two studies may be due to the sampling interval. The isolates in Tang’s study were isolated earlier, presumably closer to the original emergence event where *cfr*(C) spread on the US cattle farms mainly through an ascendant clone, followed by plasmid dissemination. The second possibility accounting for different results could be due to the different subtyping methods used to define clonality. WGS has more discriminatory power than PFGE and MLST and can provide better confirmation for clonality. The *cfr*(C) encodes resistance to several antimicrobials, including the oxazolidinone class, which is the last resort for treating MDR Gram positive bacterial infections in humans. So far, *cfr*(C) gene has not been reported in *Campylobacter* isolated from humans, FSIS regulatory samples, and retail meat, nor has *cfr*(C) gene been detected in *C. jejuni* in the US. Continued monitoring of *cfr*(C) transmission is needed to understand the spread of the gene and the MDR plasmids to different bacterial pathogens.

## Transparency declarations

None.

## Acknowledgements

We are grateful to Dr. Maureen Davidson for helpful comments and manuscript review. We also like to acknowledge Ms. Claudia Lam for handling all WGS submissions.

## Disclaimer

The views expressed in this article are those of the authors and do not necessarily reflect the official policy of the Department of Health and Human Services, the U.S. Food and Drug Administration, and Centers for Disease Control and Prevention or the U.S. Government. Mention of trade names or commercial products in this publication is solely for the purpose of providing specific information and does not imply recommendation or endorsement by the U.S. Department of Agriculture or Food and Drug Administration.

## References

1. Zhao S, Young SR, Tong E, Abbott JW, Womack N, Friedman SL, McDermott PF: Antimicrobial resistance of Campylobacter isolates from retail meat in the United States between 2002 and 2007. Appl Environ Microbiol 2010, 76(24):7949–7956.

2. Scallan E, Griffin PM, Angulo FJ, Tauxe RV, Hoekstra RM: Foodborne illness acquired in the United States--unspecified agents. Emerg Infect Dis 2011, 17(1):16–22.

3. Luangtongkum T, Jeon B, Han J, Plummer P, Logue CM, Zhang Q: Antibiotic resistance in Campylobacter: emergence, transmission and persistence. Future Microbiol 2009, 4(2):189–200.

4. Chen X, Naren GW, Wu CM, Wang Y, Dai L, Xia LN, Luo PJ, Zhang Q, Shen JZ: Prevalence and antimicrobial resistance of Campylobacter isolates in broilers from China. Vet Microbiol 2010, 144(1-2):133–139.

5. Schwarz S, Kehrenberg C, Doublet B, Cloeckaert A: Molecular basis of bacterial resistance to chloramphenicol and florfenicol. FEMS Microbiol Rev 2004, 28(5):519–542.

6. White DG, Piddock LJ, Maurer JJ, Zhao S, Ricci V, Thayer SG: Characterization of fluoroquinolone resistance among veterinary isolates of avian Escherichia coli. Antimicrob Agents Chemother 2000, 44(10):2897–2899.

7. Tang Y, Dai L, Sahin O, Wu Z, Liu M, Zhang Q: Emergence of a plasmid-borne multidrug resistance gene cfr(C) in foodborne pathogen Campylobacter. J Antimicrob Chemother 2017, 72(6):1581–1588.

8. Schwarz S, Werckenthin C, Kehrenberg C: Identification of a plasmid-borne chloramphenicol-florfenicol resistance gene in Staphylococcus sciuri. Antimicrob Agents Chemother 2000, 44(9):2530–2533.

9. Morroni G, Brenciani A, Antonelli A, D’Andrea MM, Di Pilato V, Fioriti S, Mingoia M, Vignaroli C, Cirioni O, Biavasco F et a: Characterization of a Multiresistance Plasmid Carrying the optrA and cfr Resistance Genes From an Enterococcus faecium Clinical Isolate. Front Microbiol 2018, 9:2189.

10. Hansen LH, Vester B: A cfr-like gene from Clostridium difficile confers multiple antibiotic resistance by the same mechanism as the cfr gene. Antimicrob Agents Chemother 2015, 59(9):5841–5843.

11. Deshpande LM, Ashcraft DS, Kahn HP, Pankey G, Jones RN, Farrell DJ, Mendes RE: Detection of a New cfr-Like Gene, cfr(B), in Enterococcus faecium Isolates Recovered from Human Specimens in the United States as Part of the SENTRY Antimicrobial Surveillance Program. Antimicrob Agents Chemother 2015, 59(10):6256–6261.

12. Long KS, Poehlsgaard J, Kehrenberg C, Schwarz S, Vester B: The Cfr rRNA methyltransferase confers resistance to Phenicols, Lincosamides, Oxazolidinones, Pleuromutilins, and Streptogramin A antibiotics. Antimicrob Agents Chemother 2006, 50(7):2500–2505.

13. National Antimicrobial Resistance Monitoring System Retail Meat Annual Report, 2014 [http://www.fda.gov/AnimalVeterinary/SafetyHealth/AntimicrobialResistance/NationalAntimicrobialResistanceMonitoringSystem/ucm334828.htm]

14. Zhao S, Tyson GH, Chen Y, Li C, Mukherjee S, Young S, Lam C, Folster JP, Whichard JM, McDermott PF: Whole-Genome Sequencing Analysis Accurately Predicts Antimicrobial Resistance Phenotypes in Campylobacter spp. Appl Environ Microbiol 2016, 82(2):459–466.

15. Koboldt DC, Zhang Q, Larson DE, Shen D, McLellan MD, Lin L, Miller CA, Mardis ER, Ding L, Wilson RK: VarScan 2: somatic mutation and copy number alteration discovery in cancer by exome sequencing. Genome Res 2012, 22(3):568–576.

16. Lee TH, Guo H, Wang X, Kim C, Paterson AH: SNPhylo: a pipeline to construct a phylogenetic tree from huge SNP data. BMC Genomics 2014, 15:162.

17. Chen Y, Mukherjee S, Hoffmann M, Kotewicz ML, Young S, Abbott J, Luo Y, Davidson MK, Allard M, McDermott P et al: Whole-genome sequencing of gentamicin-resistant Campylobacter coli isolated from U.S. retail meats reveals novel plasmid-mediated aminoglycoside resistance genes. Antimicrob Agents Chemother 2013, 57(11):5398–5405.

18. Wang G, Clark CG, Taylor TM, Pucknell C, Barton C, Price L, Woodward DL, Rodgers FG: Colony multiplex PCR assay for identification and differentiation of Campylobacter jejuni, C. coli, C. lari, C. upsaliensis, and C. fetus subsp. fetus. J Clin Microbiol 2002, 40(12):4744–4747.

19. Zhao S, Mukherjee S, Li C, Jones SB, Young S, McDermott PF: Cloning and Expression of Novel Aminoglycoside Phosphotransferase Genes from Campylobacter and Their Role in the Resistance to Six Aminoglycosides. Antimicrob Agents Chemother 2018, 62(1).

20. Zhao S, Mukherjee S, Chen Y, Li C, Young S, Warren M, Abbott J, Friedman S, Kabera C, Karlsson M et al. Novel gentamicin resistance genes in Campylobacter isolated from humans and retail meats in the USA. J Antimicrob Chemother 2015, 70(5):1314–1321.

21. CLSI: Performance Standards for Antimicrobial Susceptibility Testing. 26th ed. CLSI supplement M100-S26. Clinical and Laboratory Standards Institute, Wayne, PA 2016.

22. Batchelor RA, Pearson BM, Friis LM, Guerry P, Wells JM: Nucleotide sequences and comparison of two large conjugative plasmids from different Campylobacter species. Microbiology 2004, 150(Pt 10):3507–3517.

23. Bacon DJ, Alm RA, Burr DH, Hu L, Kopecko DJ, Ewing CP, Trust TJ, Guerry P: Involvement of a plasmid in virulence of Campylobacter jejuni 81-176. Infect Immun 2000, 68(8):4384–4390.

24. Bacon DJ, Alm RA, Hu L, Hickey TE, Ewing CP, Batchelor RA, Trust TJ, Guerry P: DNA sequence and mutational analyses of the pVir plasmid of Campylobacter jejuni 81-176. Infect Immun 2002, 70(11):6242–6250.

25. Ward DV, Draper O, Zupan JR, Zambryski PC: Peptide linkage mapping of the Agrobacterium tumefaciens vir-encoded type IV secretion system reveals protein subassemblies. Proc Natl Acad Sci U S A 2002, 99(17):11493–11500.

26. Fronzes R, Christie PJ, Waksman G: The structural biology of type IV secretion systems. Nat Rev Microbiol 2009, 7(10):703–714.

27. Alfredson DA, Korolik V: Antibiotic resistance and resistance mechanisms in Campylobacter jejuni and Campylobacter coli. FEMS Microbiol Lett 2007, 277(2):123–132.

28. Whitehouse CA, Young S, Li C, Hsu CH, Martin G, Zhao S: Use of whole-genome sequencing for Campylobacter surveillance from NARMS retail poultry in the United States in 2015. Food Microbiol 2018, 73:122–128.

